# Decoding the Epigenetic Landscape: Insights into 5mC and 5hmC Patterns in Mouse Cortical Cell Types

**DOI:** 10.1101/2024.07.06.602342

**Authors:** Xiaoran Wei, Jiangtao Li, Zuolin Cheng, Songtao Wei, Guoqiang Yu, Michelle L Olsen

## Abstract

The DNA modifications, 5-methylcytosine (5mC) and 5-hydroxymethylcytosine (5hmC), represent powerful epigenetic regulators of temporal and spatial gene expression. Yet, how the cooperation of these genome-wide, epigenetic marks determine unique transcriptional signatures across different brain cell populations is unclear. Here we applied Nanopore sequencing of native DNA to obtain a complete, genome-wide, single-base resolution atlas of 5mC and 5hmC modifications in neurons, astrocytes and microglia in the mouse cortex (99% genome coverage, 40 million CpG sites). In tandem with RNA sequencing, analysis of 5mC and 5hmC patterns across cell types reveals astrocytes drive uniquely high brain 5hmC levels and support two decades of research regarding methylation patterns, gene expression and alternative splicing, benchmarking this resource. As such, we provide the most comprehensive DNA methylation data in mouse brain as an interactive, online tool (**<u>NAM-Me</u>**, https://olsenlab.shinyapps.io/NAMME/) to serve as a **resource** dataset for those interested in the methylome landscape.

## INTRODUCTION

Epigenomic modifications are critical regulators of cell-type and tissue-specific, spatial and temporal gene expression across the lifespan of an organism^1–3^. Considered an impactful epigenetic regulator of gene transcription, DNA methylation, which is mediated by a family of three DNA methyltransferases (DNMT1, DNMT3A, DNMT3B)^4^, occurs predominantly at CpG dinucleotides, and involves the addition of a methyl group to the 5th carbon atom of cytosine (5mC). In brain, 5mC methylation has been extensively linked to brain development^5–7^, differentiation and maturation of neuronal cell types^8–10^, synaptic plasticity^11–15^ and learning and memory^16,17^. Moreover, alterations in brain DNA methylation are implicated in neurodevelopmental disorders^18–21^, neurodegenerative^22–24^, and neuropsychiatric disease^25–29^ and cognitive dysfunction^30–32^. 5-hydroxymethylcytosine (5hmC), an intermediate, oxidized form of 5mC represents a second abundant epigenetic mark in brain, demonstrating levels ten-fold higher in brain than observed in peripheral tissues^1,33–35^. To date, it is not known if all CNS cell types express higher 5hMC levels or if a single neural cell population drives high CNS 5hmC expression. 5hmC is mediated by ten–eleven translocation (TET) enzymes in the DNA demethylation process^36^ and serves as both a transient intermediate of DNA methylation or a stable epigenetic mark with broad genomic coverage^37–39^.

Decades of research indicate 5mC and 5hmC are powerful regulators of gene expression. 5mC expression in the promoter region of genes inhibits gene expression while 5mC in the gene body is controversial with mixed reports regarding 5mC correlates with higher gene expression^1,40^. The controversy may be due, in part, to traditional bisulfite conversion methods that lack discrimination between 5mC and 5hmC modifications. The relationship between 5hmC and gene expression is complex and context dependent. For example, gene body 5hmC positively correlates with gene expression in human GABAergic and glutamatergic neurons^41^. Similarly, in mouse embryonic stem cells, 5hmC in the promoter and gene body positively correlate with gene expression^42^ while in this same study, the level of 5hmC showed no relationship with gene expression in neural progenitor cells. In addition to regulating gene expression, 5mC expression is implicated with exon usage^43,44^, in part by orchestrating the recruitment of transcription factors as readers and effectors of DNA methylation^45^. Less is understood regarding 5hmC in exon usage, yet, it was recently demonstrated that 5hmC peaks are present at the 5’ splice site of exon-intron boundaries, suggesting 5hmC modification may serve to regulate alternative splicing^46,47^.

Cell type-specific CNS methylation profiles have been performed in human postmortem tissue^41,48,49^. Yet, neuroscience pre-clinical research is most commonly performed in murine models, necessitating an understanding of methylation patterns in unique murine CNS cell types. Here, examination of 5mC expression has revealed NeuN^+^ cells isolated from frontal cortex are more highly methylated than NeuN^-^ cells^50^. More recently, Liu et al provided a detailed single cell 5mC atlas, covering 50% of mouse genome, across 45 brain regions, revealing the DNA 5mC methylation landscape predicts single neuron cell-type identity and brain area spatial location^51^. While the vast majority of studies have focused on 5mC methylation patterns, recent technical approaches enabling discrimination between 5mC and 5hmC^52–55^, have enabled more systematic analysis of CNS 5hmC modifications. For example, Fabyanic et al used an approach termed joint single-nucleus (hydroxy)methylcytosine sequencing (Joint-snhmC-seq), which enabled simultaneous profiling 5mC and 5hmC of at single cell resolution. This approach allowed for 5mC and 5hmC methylation mapping of approximately one million CpG sites^55^ across the genome in individual cells, but due to low sequencing depth, provides low genomic coverage.

In the current study, we applied Nanopore long-read sequencing to native DNA isolated from neurons, astrocytes and microglial cells in the mouse cortex to obtain quantitative, single-base resolution identification of 5mC and 5hmC levels in each sample. This approach enabled comprehensive coverage (99% genome coverage, 40 million CpG sites) of the entire mouse genome. These results were integrated with data obtained from cell-type specific RNA sequencing, enabling a comprehensive assessment regarding the relationship between DNA 5mC and 5hmC modification patterns. Given the intense interest in DNA methylation as a primary driver of temporal and spatial gene expression in unique CNS cell populations we used these data to generate an online tool (**<u>NAM-Me</u>**, Neuronal, Astrocyte, Microglia Methylome) to serve as a benchmark for users interested the methylome landscape in preclinical murine models in the context of health and disease.

## RESULTS

### Single base 5mC and 5hmC profiling of neurons, astrocytes and microglia

To comprehensively evaluate 5mC and 5hmC level in unique CNS cell populations, we performed Nanopore long-read sequencing on DNA obtained from enriched cortical neurons, astrocytes and microglia. Nanopore sequencing was chosen as it allows for long-read, direct sequencing of native DNA^56^, provides more even genomic coverage^56^, and demonstrates lower GC bias relative to traditional approaches^57,58^. Additionally, long-read sequencing approaches including Nanopore and PacBio circumvent PCR amplification bias of chemically modified DNA, including bisulfite sequencing which can damage DNA^57,59^. Previous studies have demonstrated high 5mC and 5hmC modified base calling accuracy using Nanopore long read sequencing^60,61^. For this study, high quality, high molecular weight DNA (DIN 9.8 to 9.9, **Supplementary Figure 1A)** was extracted from cortical neurons, astrocytes and microglia cell populations using a magnetic bead isolation approach^62–65^. Each sample, per cell type was sequenced to an average depth of 13-22X, with a median read length of approximately 10,000 bp (**Supplemental Figure 1B, C**). During sequencing, as each base passes through the nanopore, it produces an electric signal (or squiggle), with unique squiggles for each base or each modified base. Subsequent base calling enables identification of each base or modified base, including 5mC and 5hmC modification. Using this approach, we consistently evaluated over 99% of all CpG sites in the mouse genome. Modified CpG sites, 5mC and 5hmC sites, were defined as any CpG site with modification levels greater than 10%^66,67^. Analysis at single base resolution revealed approximately 41 million 5mC sites in neurons, astrocytes and microglia met the 10% or greater 5mC level threshold **(Figure 1A).** Our analysis revealed significantly different 5mC distribution among all three cell types. Microglia demonstrate significantly more CpG sites with higher 5mC level than when compared to neurons and astrocytes (Kruskal-Wallis test, *p* < 2.2e-16), with over half of the microglia CpG sites demonstrating 80% or higher 5mC level (**Figure 1A, B**). Neurons also demonstrated higher levels of 5mC modified bases compared to astrocytes (Kolmogorov-Smirnov Test, *p* < 2.2e-16), with approximately 35% and 30% of their genomes highly 5mC modified (> 80% 5mC level, **Figure 1B**). Each 5mC site, in each cell population, was then mapped into one of six different genomic regions; the promoter (1Kb upstream of the transcription start site), 5’UTR, CDS, introns, 3’UTR, and intergenic regions. This analysis indicated a 5mC odds ratio (compared to CpG sites) of around 1.0 for all genomic locations, with notable exceptions including gene promoters and 5’ UTR regions which demonstrated the lowest probability of 5mC modifications **(Figure 1C)**.

**Figure 1:**
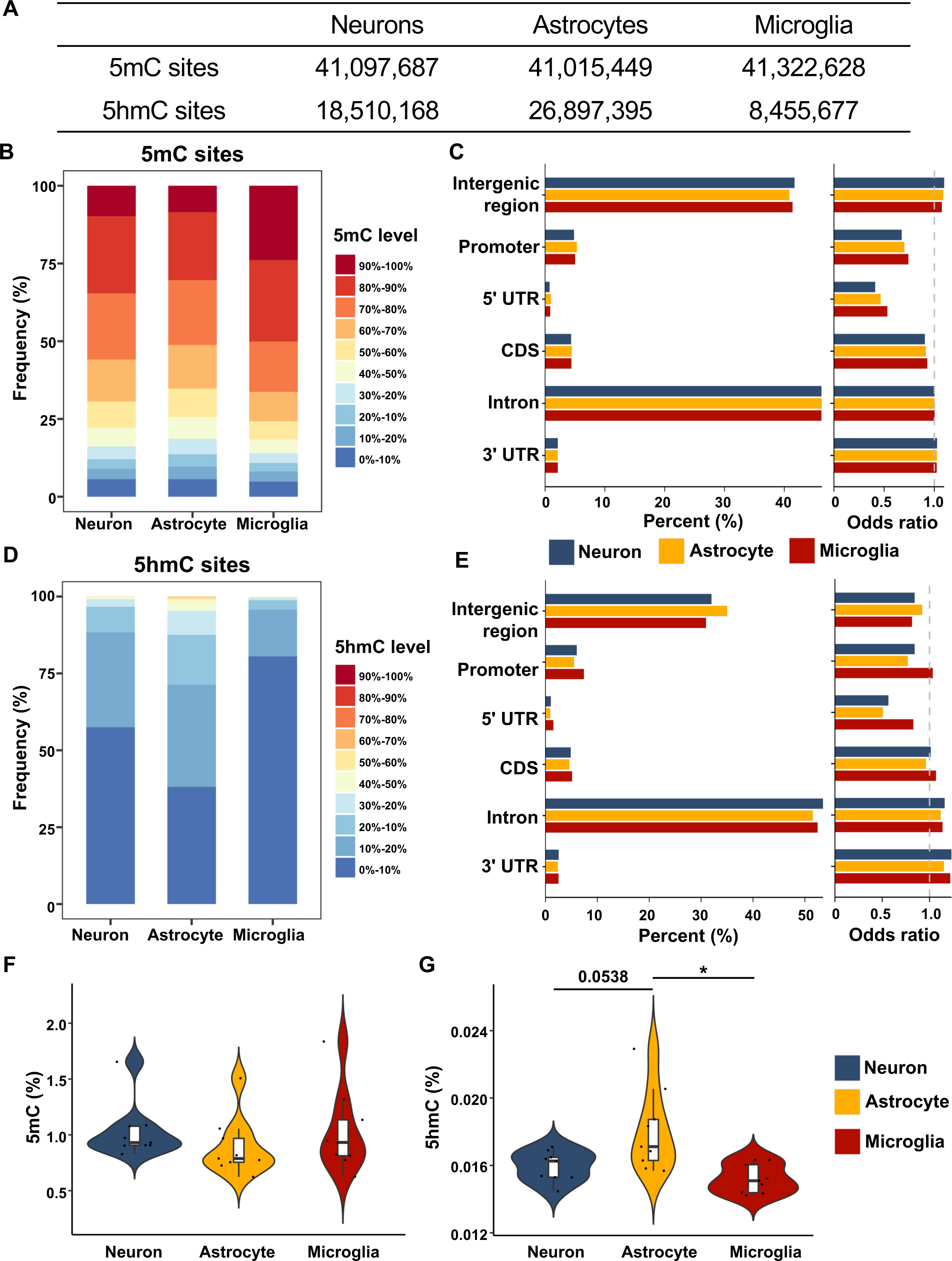
Single base 5mC and 5hmC profiling of astrocyte, neuron, and microglia. **A,** Table of the identified 5mC and 5hmC sites (5mC/5hmC level > 10%) in neurons, astrocytes and microglia. **B and D**, Distribution of 5mC level (**B**) and 5hmC level(**D**) across all identified CpG sites in neurons, astrocytes and microglia. About 95% of CpG sites have 5mC modification (5mC level > 10%) in all three cell types while 62% of CpG sites show 5hmC modification (5hmC level > 10%) in astrocytes, 42% in neurons and only 19% in microglia. 3 biological replicates are involved in each cell type. Distribution of both 5mC and 5hmC are significantly different across cell types, Kruskal-Wallis test, *p* < 2.2e-16. **C and E**, Genomic distribution of 5mC (**C**) and 5hmC (**E**) sites (5mC/5hmC level > 10%). Odds ratio was calculated based on the genomic distribution of CpG sites in mouse genome. **F-G,** violin plot of the global 5mC (**F**) and 5hmC (**G**) level in neurons, astrocytes and microglia (n = 9, ANNOVA test).

We next quantified the occurrence of 5hmC modifications, at single base resolution, for each cell type using the same 10% threshold. This analysis revealed striking differences in the level of 5hmC modified bases in each cell type. Astrocytes demonstrated significantly more CpG sites with higher levels of 5hmC modifications compared to neurons or microglia (Kruskal-Wallis test, *p* < 2.2e-16), with 62% of CpG sites greater than 10% 5hmC level (**Figure 1 E)**. Accordingly, microglia which demonstrated the highest levels of 5mC modifications, showed relatively low levels of 5hmC modified bases (<20% 5hmC level, **Figure 1E)**. These data suggest that the reported high level of 5hmC modification observed in the central nervous system^1,33–35^ is mediated, in part, by astrocyte 5hmC modified bases and also reveals heterogenous 5mC and 5hmC patterns in astrocytes and microglia. Mapping 5hmC modified bases to genomic location, we identified 5hmC modified bases overrepresented in both introns and the 3’ UTR. The lowest odds ratio was again observed in the 5’ UTR **(Figure 1E)**.

To validate the above nanopore sequencing data, we performed ELISA assays to assess the global levels of 5mC and 5hmC in all three cell types **(Figure 1F, G)**. Using this approach we were unable to identify significant differences in 5mC levels among the cell types, yet, astrocytes exhibited a trend towards the lowest 5mC levels, while microglia displayed a trend towards the highest levels **(Figure 1F)**. Regarding 5hmC levels, astrocytes showed significantly higher levels than microglia and a trend towards higher levels compared to neurons (*p value* = 0.0538) **(Figure 1G),** consistent with the trends observed in the nanopore sequencing data. Together, these data reinforce 5mC as the predominant DNA modification, and show astrocytes, in particular, contribute to high CNS 5hmC levels.

Long read, native DNA Nanopore sequencing, enables accurate sequencing and mapping of repetitive elements and CG-rich regions. Thus, we investigated the 5mC patterns in repetitive elements, CpG islands and CpG shores **(Figure 2A-C)**. DNA 5mC methylation is typically considered to be stable and high in repetitive elements^68,69^, which constitute over 50% of human genome^70,71^. Further, DNA methylation is often low in the CpG rich regions in gene promoters^72^. CpG shores, which are 2kb away from the CpG islands, have also been demonstrated to show low methylation levels^73^. Here, we found 5mC levels are significantly different among each region in all three cell types (Kruskal-Wallis test, *p* < 2.2e-16). We confirmed that compared to the whole genome, a greater percentage of CpG sites demonstrate high 5mC level in repetitive elements **(Figure 2A).** We identified that over 75% repetitive elements in all three cell populations demonstrated methylation levels at or above 60-70%. In contrast, and as expected, CpG island and CpG shores demonstrated significantly lower levels of 5mC methylation **(Figure 2B, C)**. 5hmC methylation levels also show significant regional differences (Kruskal-Wallis test, *p* < 2.2e-16) across each of these regions, where CpG sites in repetitive elements and CpG islands have fewer CpG sites with high 5hmC level when compared to the whole genome **(Figure 2D, E)**, while CpG shores demonstrate a greater number of CpG sites with high 5hmC when compared to the remainder of the genome **(Figure 2F)**.

**Figure 2:**
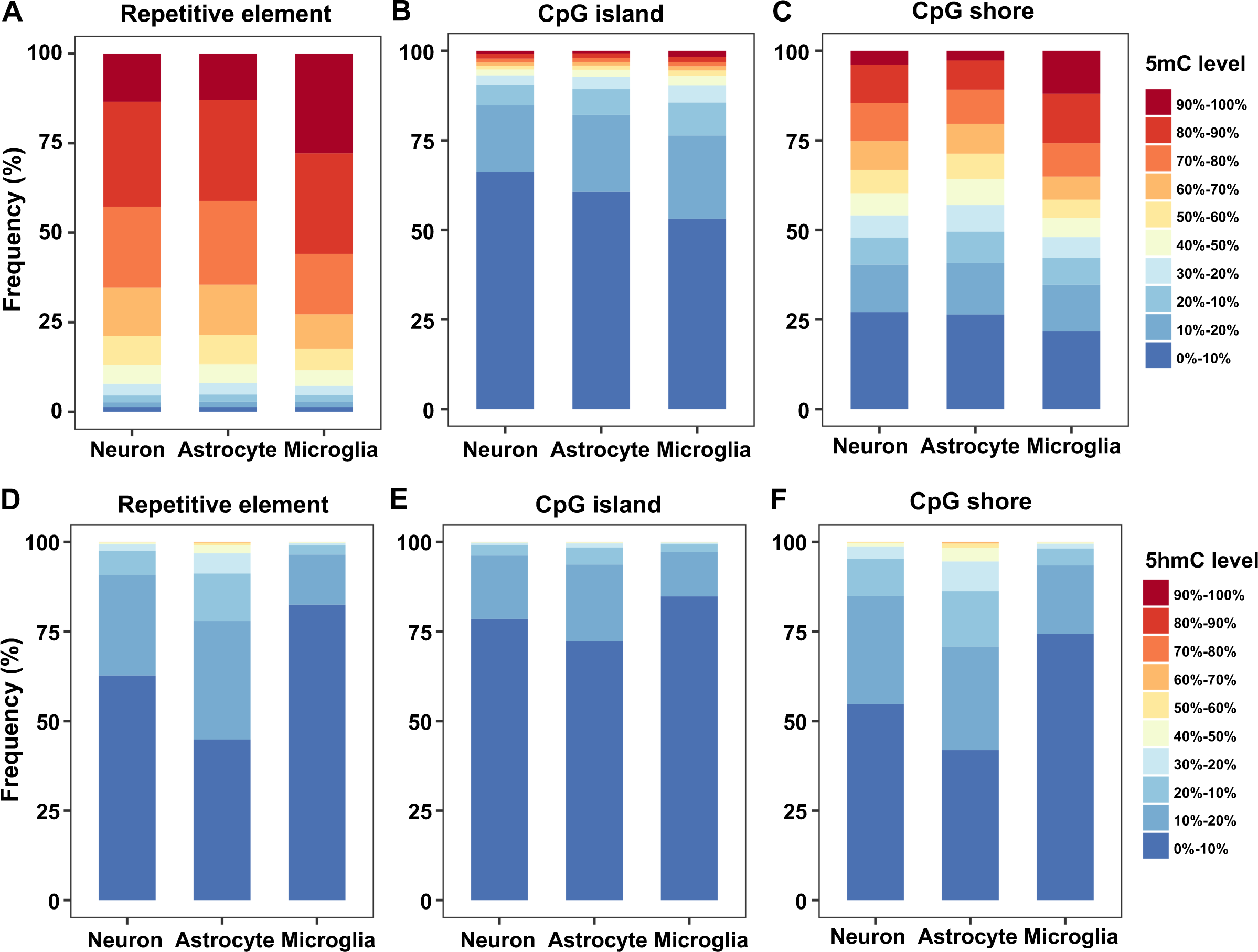
5mC and 5hmC level in repetitive element, CpG island and CpG shore. **A-C**, Distribution of 5mC level of across CpG sites in repetitive element (**A**), CpG island (**B**) and CpG shore (**C**). **D-F**, Distribution of 5hmC level of all CpG sites in repetitive element (**D**), CpG island (**E**) and CpG shore (**F**).

### Differential 5mC and 5hmC profiles across CNS cell populations

Given the methylation differences observed in neurons, astrocytes and microglia, we next sought to evaluate differential 5mC and 5hmC across all three cell populations. To this end, we identified differentially methylated sites (DMSs) and differentially hydroxymethylated sites (DhMSs), defined as any single base with a 5mC/5hmC level difference greater than 10% (FDR < 0.01)^74,75^ between two unique cell populations. When comparing astrocytes to neurons, we identified over 4 million DMSs, the majority of which (78.6%) are hypermethylated in neurons **(Figure 3A, B)**. Manhattan plots demonstrate the distribution pattern across each chromosome and indicates over 99.8% of DMSs between astrocytes and neurons is below 50% **(Figure 3B)**. We next applied the same criteria to identify DhMSs (> 10% 5hmC level difference, FDR < 0.01). These analyses reveal the 5hmC bias in astrocytes relative to neurons, with > 99% of the 1,772,951 DhMSs hypermethylated in astrocytes **(Figure 3C, D)**. DMS and DhMS analysis was also performed for astrocytes vs. microglia and neurons vs. microglia (**Supplemental Figure 2**). We observed that Y-chromosome DMS and DhMS density was low when compared to other chromosomes, which may relate to homogenous Y-chromosome protein coding gene expression across each cell type (not shown). Each DMS and DhMS was again annotated to the same six genomic regions as above **(Figure 3E, F)** and then collapsed into differentially methylated regions (DMRs) and differentially hydroxymethylated regions (DhMRs) by clustering the DMSs or DhMSs (**Supplementary** Figure 3**, 4**). Together, these findings support findings that across the entire genome, astrocytes show high levels of 5hmC methylation relative to neurons and microglia.

**Figure 3:**
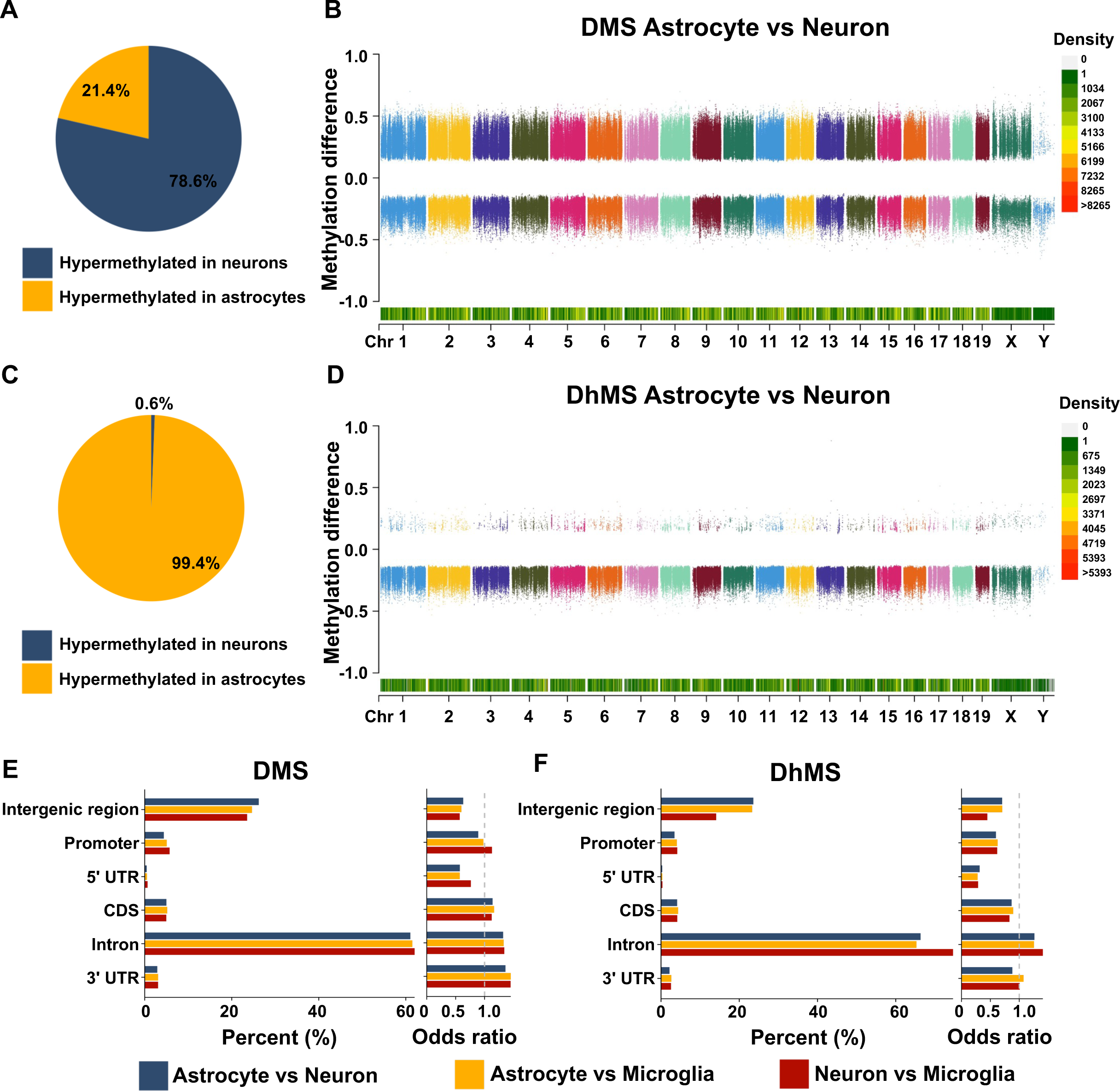
5mC and 5hmC level difference across cell types. **A**, Distribution of hypermethylated DMSs between astrocytes and neurons. 4,083,859 DMSs (methylation difference > 0.1 and FDR < 0.01) were identified between astrocytes and neurons. 78.6% of all DMSs are hypermethylated in neurons (3,210,519 DMSs) while 21.4% are hypermethylated in astrocytes (873,335 DMSs). **B,** DMSs between astrocytes and neurons. All the CpG sites are plotted based on their chromosome location and modification level difference between neurons and astrocytes (astrocytes served as the control). **C**, Distribution of hypermethylated DhMSs between astrocytes and neurons. 1,772,951 DhMSs (methylation difference > 0.1 and FDR < 0.01) were identified and 99.4% of the DhMSs are hypermethylated in astrocytes (1,762,382 DhMSs). **D**, DhMSs between astrocytes and neurons. All CpG sites are plotted based on their chromosome location and modification level difference between neurons and astrocytes (astrocytes served as the control). **E-F**, Genomic distribution of DMSs (**E**) and DhMSs (**F**). The odds ratio was calculated based on the genomic distribution of 5mC or 5hmC sites in mouse genome.

### 5mC and 5hmC DNA modification and cell type specific gene expression

To examine the relationship between 5mC and 5hmC modification and gene expression, we next integrated nanopore data with coding gene expression data across each cell population. For these studies, RNA was isolated from cortical neurons, astrocytes and microglia using a magnetic bead approach, from age- and sex-matched littermates from the Nanopore studies above. The relative purity of each individual cell population as assessed by gene expression correlation, frequency of relative gene expression, and quantitative PCR are shown in **Supplemental Figure 1D-G**. 5mC levels along the promoter region and extending into the first exon, were plotted for each cell population. These data revealed 5mC level was at its lowest, or reached a valley, immediately before the transcription start site (TSS) **(Figure 4A).** While generally, 5hmC levels are lower across the entire genome, we observed a similar valley for 5hmC modification immediately preceding the TSS **(Figure 4B).** To investigate the relationship between promoter region 5mC/5hmC level and gene expression, we calculated the weighted promoter modification level for either 5mC or 5hmC and correlated to gene expression level for each cell type. Here, weighted modification level indicates the averaged level across all CpG sites in the promoter region. As expected, this analysis revealed a consistent negative correlation between 5mC promoter level and gene expression **(Figure 4C)**, while no clear relationship between 5hmC promoter level and gene expression was observed (|r| < 0.15) **(Figure 4D)**. A weak negative relationship was also observed for 5mC level and gene expression level in the gene body **(Figure 4E)**. Comparing to the r-value between promoter 5mC level with gene expression, 5mC on the gene body showed a weaker corelation with gene expression. A positive relationship between 5hmC level and gene expression **(Figure 4F)**. Density plots reveal 5mC and 5hmC levels are markedly higher in the gene body relative to the promoter region in all three cell types **(Figure 4G, H)**.

**Figure 4:**
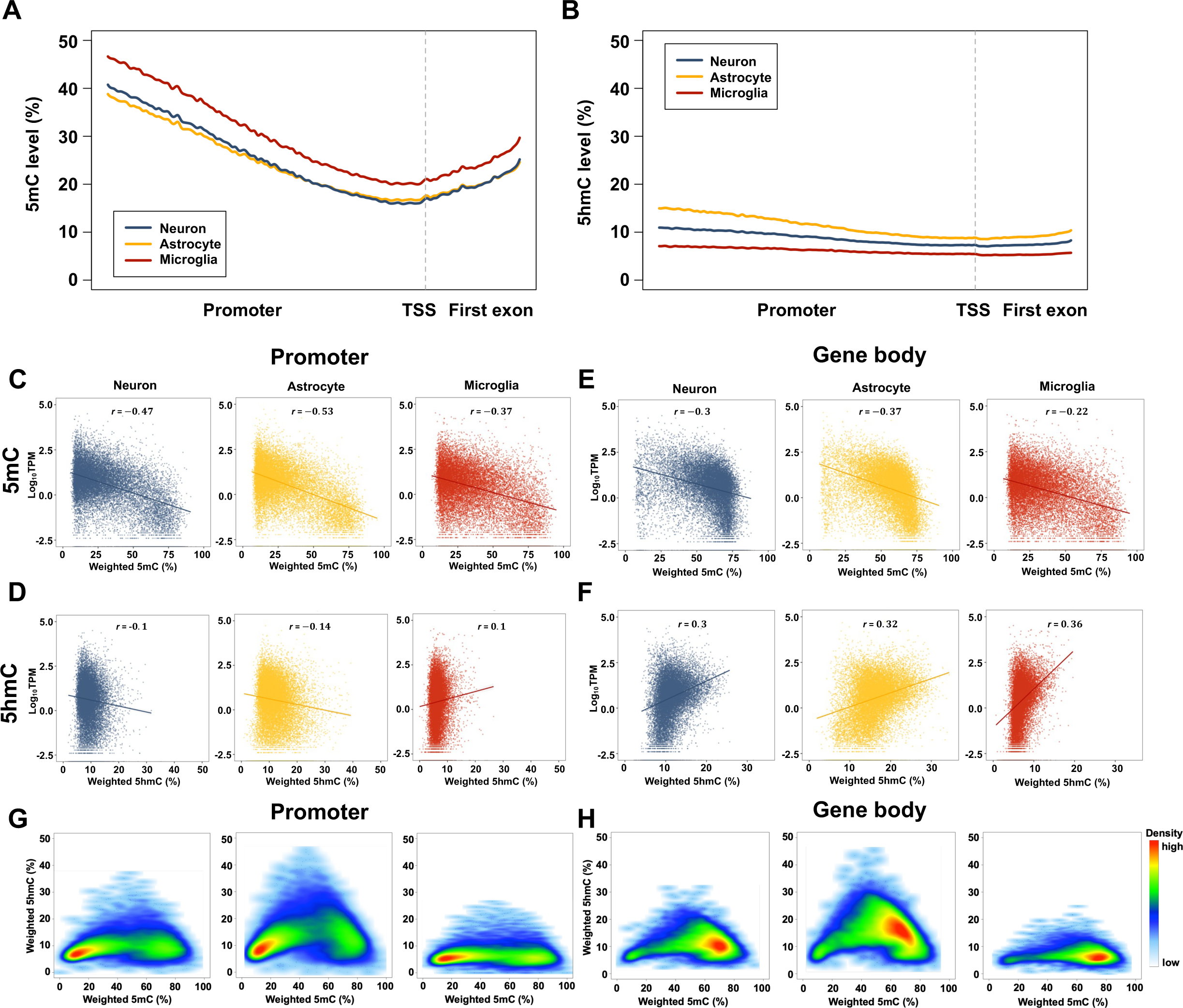
DNA methylation and gene expression. **A-B**, Distribution of 5mC (**A**) and 5hmC (**B**) levels among promoter and first exon. **C-D**, Correlation of expression levels of gene expression level and weighted 5mC (**C**) or 5hmC (**D**) level of each gene’s promoter region in neurons, astrocytes and microglia. Each point represents one gene. The lines are fitted with a linear regression. *p* < 0.001 for all associations. **E-F**, Correlation of expression levels of gene expression level and weighted 5mC (**E**) or 5hmC (**F**) modification level of each gene’s gene body in neurons, astrocytes and microglia. *p* < 0.001 for all associations. **G-H**, Density plot of correlation of weighted 5mC level and 5hmC level in the promoter region (**G**) and gene body (**H**) in neurons, astrocytes and microglia.

The above analysis represents general trends across all protein coding genes and their relative expression levels in relationship to 5mC and 5hmC modifications. In the next series of analyses, we correlate differential DNA methylation to genes that are exclusively expressed in one cell population. Here cell-type specific genes were defined as genes that 1) met standard statistical criteria for differential expression, 2) exhibited a 20-fold higher expression or greater in the target cells compared to the other two cell types, and 3) had a TPM > 50. We also manually evaluated all candidate genes on Dropviz^76^ to validate gene expression is highest in the target cells. Using this approach 24 astrocyte and 100 microglia specific genes were identified, 78 of which have been identified as core human microglial genes^77^ (**Figure 5A, B, Supplementary Table 1)**. No neuronal genes met these criteria. The neuronal gene *Tac2* demonstrated > 20-fold expression in neurons, but had an average TPM of 37.4. Related to astrocytes and microglia, several of the genes we identified using our defined criteria have been described as cell-type markers including, astrocyte *Chrdl1*^78^, *Sox9*^79^ and *Aldh1l1*^80^ as well as microglial *Itgam*^81,82^, *Trem2*^83^ and *Tmem119*^84^. Across all 24 astrocyte specific genes, 33 DMRs were identified in the promoter when compared to neurons (**Figure 5C**), and 38 promoter DMRs were identified when compared to microglia (**Figure 5D**). Remarkably, almost all DMRs were significantly hypomethylated in astrocytes when compared to neurons (97.0%) and microglia (97.4%) **(Figure 5C, D)**. Similarly, 100% of astrocyte promoter DhMRs were hypermethylated relative to neurons and microglia **(Figure 5E, F)**. Focusing on gene body methylation we observed the vast majority of DMRs on astrocyte genes (87.8% compared to neurons and 84.8% compared to microglia) are hypomethylated in astrocytes **(Figure 5G, H)**, while 99.9% and 100% astrocyte DhMRs are hypermethylated relative to neuronal and microglial cell populations **(Figure 5I, J)**. We selected one astrocyte gene, *Aldh1l1* (an aldehyde dehydrogenase), for further evaluation of 5mC and 5hmC by mapping each DMS and DhMS within the 1000 bp promoter region and gene body (**Figure 5K**). Here, the absolute 5mC and 5hmC level at each DMS and DhMS is shown for all three cell types, with average methylation values for the entire region summarized in **Figure 5L**.

**Figure 5:**
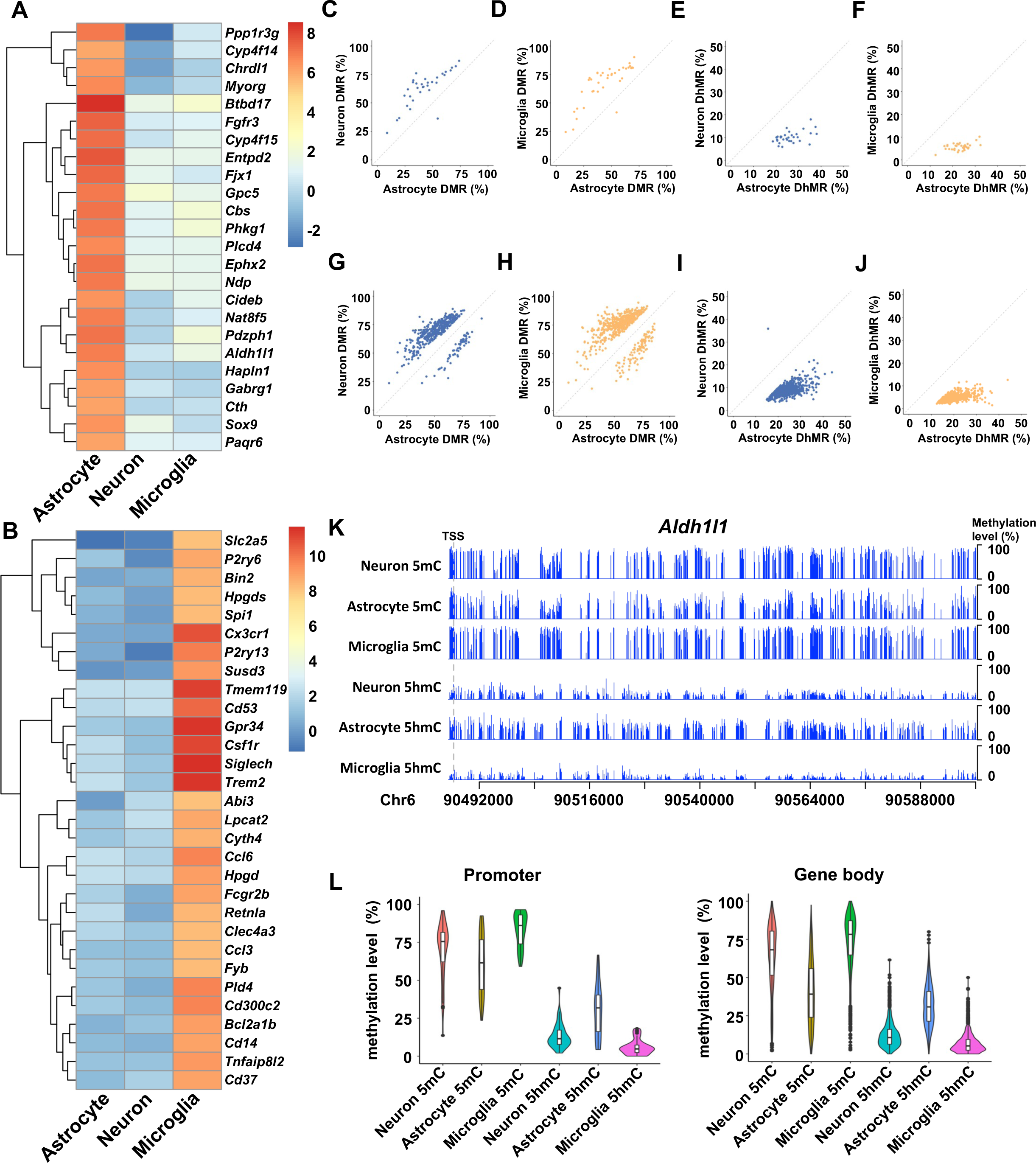
DNA methylation and cell-type specific gene expression. **A-B**, Gene expression levels of astrocyte specific genes (**A**) and microglia specific genes (top30) (**B**) in astrocytes, neurons and microglia. **C-D**, 5mC level of DMRs between astrocytes and neurons (**C**) and astrocytes and microglia (**D**) in the promoter region of astrocyte specific genes. **E-F**, Modification level of DhMRs between astrocytes and neurons (**E**) and astrocytes and microglia (**F**) in the promoter region of astrocyte specific genes. **G-H**, 5mC level of DMRs between astrocytes and neurons (**G**) and astrocytes and microglia (**H**) in the gene body region of astrocyte specific genes. **I-J**, 5hmC level of DhMRs between astrocytes and neurons (**I**) and astrocytes and microglia (**J**) in the gene body region of astrocyte specific genes. **K**, Methylation status in Aldh1l1. The gray dash line represents TSS. **L**, Violin plot of 5mC and 5hmC levels in the promoter and gene body of *Aldh1l1* in different cell types.

A similar analysis was also performed for microglia specific genes. Generally, we observed a weaker relationship between microglial gene expression and 5mC promoter level with approximately 73% of microglia DMSs hypomethylated relative to astrocytes and neurons **(Supplementary Figure 5A, B)**. Little discrimination was observed for 5mC gene body modification or 5hmC promoter gene body modification (**Supplementary Figure 5C-H)**. We performed a similar analysis for the gene *Tac2*, however only one DMR was identified in the promoter region when compared with astrocytes, and was hypermethylated in neurons (data not shown). To identify DNA methylation patterns in neuron specific genes, we decreased the criteria to a greater 10-fold change compared to astrocytes and microglia and decreased TPM to > 20 TPM in neurons. Five such neuron specific genes were identified (**Supplementary Figure 5I)** but did not contain sufficient DMRs or DhMRs to compare to astrocytes or microglia (data not shown). Here, the various subtypes of neurons collected together, may account for our inability to identify neuronal specific genes.

Accordingly, when focusing on cell type specific gene expression, these results support enhanced 5mC modifications in the promoter region is a critical mechanism for gene expression inhibition in astrocytes and microglia. Additionally, these results reveal, at least in astrocytes both promotor and gene body 5hmC modification may cooperate to promote astrocyte specific gene expression.

### 5mC and 5hmC DNA methylation and cell type-specific alternative splicing

Alternative splicing contributes to transcript and subsequent proteomic diversity. Alternative splicing is mediated by complex molecular coordination, including cis-acting sequence signals, RNA binding proteins, 3D chromatin structure, and epigenetic modifications, including DNA methylation ^85–87^. This comprehensive genomic 5mC and 5hmC dataset and complimentary protein coding gene expression dataset provide unique opportunity to further explore the relationship between methylation and exon usage. Comparing neurons to astrocytes we identified 4,436 differentially used exons (log_2_FC > 1, adjusted p-value < 0.05) (**Figure 6A**). For the comparison of astrocytes to microglia and neurons to microglia, we identified 10,285 and 5909 differentially used exons, respectively (**Figure 6B, C**). 5mC and 5hmC level along exon and exon-intron boundaries (250 bp upstream and downstream) was plotted for all constitutive exons and alternative exons for each cell type (**Figure 6D-I**). These analyses revealed a clear pattern of higher 5mC and 5hmC level in exons relative to introns and a characteristic dip in methylation at the intron-exon boundaries. Constitutive expressed exons (solid lines) also showed higher 5mC levels when compared to alternatively expressed exons in both exon and adjacent intron regions (**Figure 6D-F**). This trend was also observed for 5hmC modification (**Figure 6G-I**).

**Figure 6:**
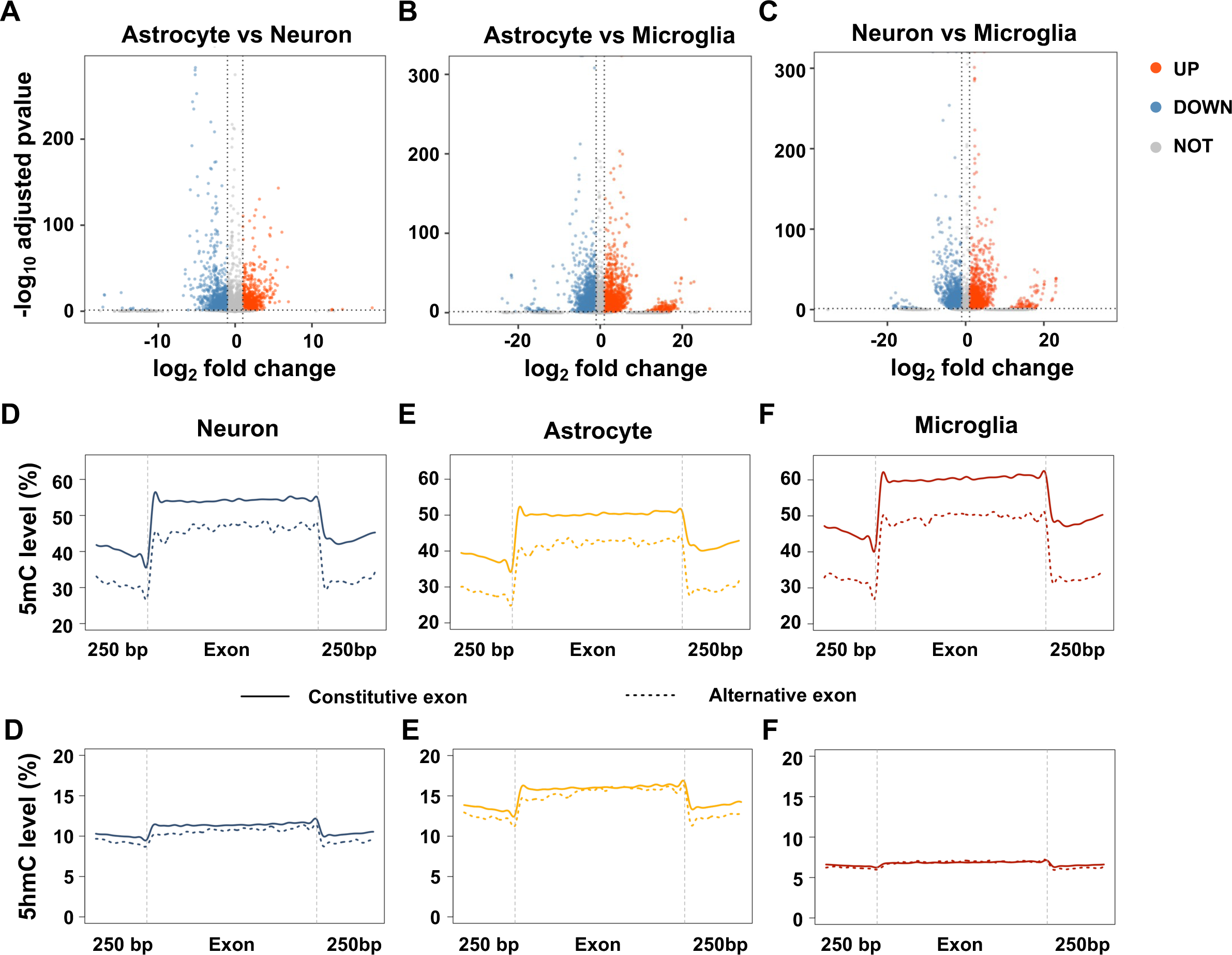
DNA methylation regulates alternative splicing process. **A-C**, Differentially used exons (Fold change > 2, adjusted p-value < 0.05) between astrocytes and neurons (**A**), astrocytes and microglia (**B**), neurons and microglia (**C**). **D-F**, Distribution of 5mC levels among constitutive and alternative exons and exon-intron boundaries in neurons (**D**) astrocytes (**E**) and microglia (**F**). The solid line represents constitutive exon and the dotted line represent alternative exon. **G-I**, Distribution of 5mC levels among constitutive (solid line) and alternative exons (dotted line) and exon-intron boundaries in neurons (**G**) astrocytes (**H**) and microglia (**I**).

## DISCUSSION

In this study, we report genome-wide 5mC and 5hmC modification of neurons, astrocytes and microglia from mouse cortex at single-base resolution using Nanopore long-read sequencing. Our studies, for the first time reveal heterogenous 5mC and 5hmC methylation patterns across microglia and astrocytes, whereby microglia, a peripherally derived CNS cell type demonstrates the highest 5mC levels; >50% of all CpG sites demonstrating greater than 80% methylation (**Figure 1B**). Our results also consistently indicate astrocytes demonstrate high 5hmC methylation, relative to neurons and microglia with approximately 60% of all CpG sites demonstrating 10% or greater 5hmC methylation, and was predictive for identifying astrocyte enriched genes. These findings indicate astrocytes contribute to high 5hmC levels repeatedly identified in brain relative to peripheral tissues^1,33–35^ and may be a mechanism whereby astrocytes, which are born later in development from a common neural progenitor pool as neurons, serve to regulate their gene expression^88^. Our results also serve to confirm multiple studies, across a decade or more of methylation research, benchmarking this data set as a resource dataset for those interested in the methylome landscape, while also presenting an opportunity for further data mining for alternative DNA modifications in future studies.

Here we quantified 5mC and 5hmC levels across 40 million CpG (99% genome coverage, approximately 20X coverage per sample, 60X coverage per cell population) sites in neurons, astrocytes and microglia, providing the most comprehensive evaluation of CNS cell-type specific 5mC and 5hmC modification in the murine cortex. Methylation sequencing of native DNA negated a need for bisulfite conversion or chemical protection of 5hmC to detect both 5mC and 5hmC in the same cell populations. In addition to extensive coverage, our study for the first time characterizes 5mC and 5hmC at each GpG site in astrocytes and microglia cell populations and demonstrates marked heterogeneity of the methylome in these two CNS cell types. Previous research in oligodendrocytes, a glial cell type, demonstrated lower levels of 5mC and 5hmC methylation relative to NeuN^+^ cells^41,48^. Here, we demonstrate microglia show high 5mC levels when compared to neurons while astrocytes have higher 5hmC level than neuron in our study. Further, our studies are in contrast to a recent publication demonstrating astrocytes have relatively low levels of 5hmC methylation utilizing Joint-snhmC-seq^55^, which performed methylation analysis in single cells, including a small number of astrocytes (28 cells). Due to the approach utilized here, enriching astrocytes across the entire cortex, we can not exclude the possibility unique astrocyte populations may demonstrate variability across their methylome, including populations with low levels of 5hmC methylation. Notably, differences in results across these two studies may also derive from reported genomic coverage (1 million vs. 40 million CpG sites) and coverage at each individual CpG site.

Here we also evaluated the relationship between cell-type specific coding gene expression and 5mC and 5hmC modifications with all coding genes expressed in a single cell population (Figure 3) and cell type specific genes (Figure 4). At the global level, our data supports a negative correlation between promoter and gene body 5mC modification and gene expression^1,40^. In contrast, no clear relationship was observed between 5hmC promoter modification and cell-type specific global gene expression. However, 5hmC in gene body demonstrated a positive relationship with gene expression in all three cell types, consistent with previous findings in human GABAergic and glutamatergic neurons and mouse embryonic stem cells^41,42^. When focusing on cell type-specific genes, we observed the relationship between 5mC/5hmC modification and gene expression was stronger in astrocytes, such that 100% of DMRs in the promoter of astrocyte showed significantly lower 5mC level relative to the other two non-expressing cell types. This result was similarly striking for 5hmC modification where DhMRs in the promoter and gene body of astrocyte specific genes were hypermethylated when compared to the 5hmC levels in the other two cell populations, suggesting in astrocytes 5mC and 5hmC modification coordinate to regulate cell type specific genes. However, this relationship was not observed in microglia. These studies are in line with previous results indicating 5hmC plays a context-dependent and cell type specific role in gene expression and in cells expressing high 5hmC levels in the promoter and gene body, this modification positively correlates with gene expression^42^.

Collectively, these data provide a rich resource and as such, we have generated an online interactive website (**NAM-Me**, Neuronal, Astrocyte, Microglia Methylome) allowing users to identify 5mC and 5hmC modification by gene, or chromosomal location, and visualize and compare differentially methylated sites or download data for any gene or gene region in each cell type. NAM-Me serves as a resource dataset for researchers interested in exploring the methylome landscape in preclinical murine models in the context of health and disease.

## Acknowledgments

We thank Dr. Song Li for sharing the PromethION to perform Nanopore sequencing. We thank Megan Naff and Jennifer Jenrette from The Genomics Sequencing Center at Virginia Tech for helping with Nanopore sequencing. We thank Dr. Georgia Hodes and Dr. Timothy Jarome for participating in the discussions related to manuscript preparation.

This work was supported by the National Institutes of Health [R01NS120746 to M.O.].

## Author contributions

X.W., J.L., and M.O. conceptualized, and conducted experiments performed data analysis, and wrote the manuscript. Z.C, S.W. and G.Y. participated in data analysis.

## MATERIAL AND METHODS

### Animals

Wild-type C57BL/6 male mice were housed and bred at Virginia Polytechnic Institute and State University. All experiments performed were approved by Virginia Polytechnic Institute and State University Animal Care and Use Committee. Animals were maintained on a reverse 12 hr light/dark cycle (lights on at 10 pm, lights off at 10 am) with food and water available ad libitum. All tissue was collected between 10 am and 2 pm. To minimize potential litter effects, animals from at least three different litters were randomly assigned to each experimental group.

### Sequential cell isolations

Cells were isolated as previously described^62–65^. Briefly, mice were anesthetized with CO2 on postnatal day 28 +/- 1 day (P28). Whole cortices were dissected in ice cold, 95% O2 bubbled ACSF (120 mM NaCl, 3.0 mM KCl, 2 mM MgCl, 0.2 mM CaCl, 26.2 mM NaHCO3, 11.1 mM glucose, 5.0 mM HEPES with 3 mM AP5, 3 mM CNQX) and single cell suspensions were acquired with an entire mouse cortex using a papain dissociation kit (Worthington Biochemical, Lakewood, NJ, USA, #LK003153). Cell suspensions were incubated with Myelin removal microbeads (Milteny Biotech, Teterow, Germany, #130-096-733) for 10 min to remove oligodendrocytes. The flow through was captured and the cell suspension was split in half to capture microglia and astrocytes or neurons as we have described^64,65^.

### Nanopore Sequencing

High molecular weight (HMW) DNA was extracted from isolated cells using the Monarch HMW DNA extraction kit (New England Biolabs, Ipswich, MA, USA, #T3050L) according to the manufacturer’s instructions. The HMW DNA was sheared to 10kb segments with g-TUBE (Covaris, Woburn, MA, USA, #520079). Libraries were prepared and sequencing adapters were attached according to Oxford’s Ligation Sequencing Kit (Oxford Nanopore Technologies, Oxford, United Kingdom, #SQK-LSK110) instructions. One to two flow cells were utilized for each library to ensure an optimal sequencing depth of around 20x coverage at each cytosine and sequenced on PromethION (Nanopore Technologies, The Genomics Sequencing Center, Virginia Tech). Three biological replicates were included for each group. Following the sequencing, raw data was stored in fast5 files.

### Nanopore Sequencing Analysis

Fast5 file raw signals were converted to nucleotide sequences, which were mapped to the mouse reference genome mm10 using Megalodon (v2.4.2). Quality control was performed with pycoQC (v2.5.2) using the sequencing summary file generated by Megalodon. Methylated CpG sites were distinguished based on the electronic signal difference and mapping result with the model provided by Remora (v0.1.2). The final output provides the position and quantitative modification level of each cytosine site for both 5mC and 5hmC modification. Only CpG sites with average coverage greater than 3 and identified in at least two libraries were used for the following analysis. Methylated sites were defined as CpG sites with modification levels over 10%. DMSs/DhMSs and DMRs/DhMRs among neurons, astrocytes and microglia were identified with the R package DSS (v2.44.0)^89^ with smoothing = T setting. Only sites with 5mC/5hmC level difference greater than 10% and FDR less than 0.01 were considered as DMSs/DhMSs. DMRs/DhMRs were defined as any region over 50bp with at least 3 CpG sites, with > 50% of CpG sites in a region demonstrating a significant 5mC/5hmC level difference (FDR <0.01) and the average 5mC/5hmC level across all CpG sites in a region should be greater than 10%^74^.

### RNA isolation and qPCR

Isolated cells were stored in Trizol (−80°C) prior to RNA isolation. RNA was isolated using the Direct-zol RNA Microprep kit (Zymo Research, Irvine, CA, USA, #R2060) according to the manufacturer’s instructions. 4ng RNA was reverse transcribed into cDNA using iScript™ Reverse Transcription Supermix (Bio-Red, Hercules, CA, USA, #1708841). Neuron, astrocyte and microglia enrichment were determined by qPCR. Here Taqman PCR master mix (Thermo Fisher Scientific, Waltham, MA, USA, #4444557) and TaqMan probes Rbfox3 for neurons, Gfap for astrocytes, Itgam for microglia and MBP for oligodendrocytes were used for this purpose, with GAPDH serving as the housekeeping gene. The ddCt method was employed to determine the relative mRNA expression levels.

### RNA sequencing

Library preparation and RNA sequencing were performed by Novogene Co using the SMART-Seq v4 PLUS Kit (Takara Bio, Kusatsu, Shiga, Japan, # R400752). Total RNA was reverse transcribed into the first-strand cDNA. Double-strand cDNA was synthesized by LD-PCR amplification, followed by purification with AMPure XP (Beckman Coulter, Brea, CA, USA, #A63882) beads and quantification with Qubit. PolyA enrichment was conducted to enrich all the coding RNA. The cDNA samples were then fragmented, end-repaired, A-tailed, and ligated with adaptors. After size selection and PCR enrichment, sequencing was performed on a NovaSeq instrument (Illumina, San Diego, CA, USA). Paired-end reads 2 x 150 bp sequencing runs were performed with an average of 30 million reads per sample. Five biological replicates were included in each group.

### RNA sequencing analysis

Bases with quality scores less than 30 and adapters were trimmed from raw sequencing reads by Trim Galore (v0.6.4). After trimming, reads with lengths greater than 30bp were mapped to mm10 by STAR (v2.7.1a) with an average mapping efficiency of 81.5%. Raw counts and normalized counts for each gene were produced by RSEM (v1.2.28). The raw counts were used to identify differentially expressed genes (DEG) using DESeq2 (v1.36.0). Only genes with an adjusted p-value less than 0.05 and at least two-fold change were considered as DEGs. Exon usage difference was performed with DEXseq (v1.44.0). Exons with an average count greater than 10 in at least one group, adjusted p-value less than 0.05, and at least two-fold change were considered as differentially used exons.

### 5mC and 5hmC ELISA

Neurons, astrocytes and microglia were isolated as described above. Genomic DNA was extracted from the cells using the Purelink Genomic DNA mini kit (Invitrogen, Waltham, MA, USA, #K182001). For each well, 100 ng of DNA was used for the 5mC ELISA (EpigenTek, Farmingdale, NY, USA, #P-1030-96) and 5hmC ELISA (EpigenTek, Farmingdale, NY, USA, #P-1032-96) and global 5mC and 5hmC in each sample was quantified according to the manufacturer’s instructions. In total, 9 individual samples were isolated and assessed from 9 WT cortices at P28.

## Data availability

Raw Nanopore Sequencing data including fast5 files and fastq files and raw RNA Sequencing data fastq files are available in GEO database with accession number GSE244256.

