## Supplemental Figures for "Decoding the Epigenetic Landscape: Insights into 5mC and 5hmC Patterns in Mouse Cortical Cell Types"

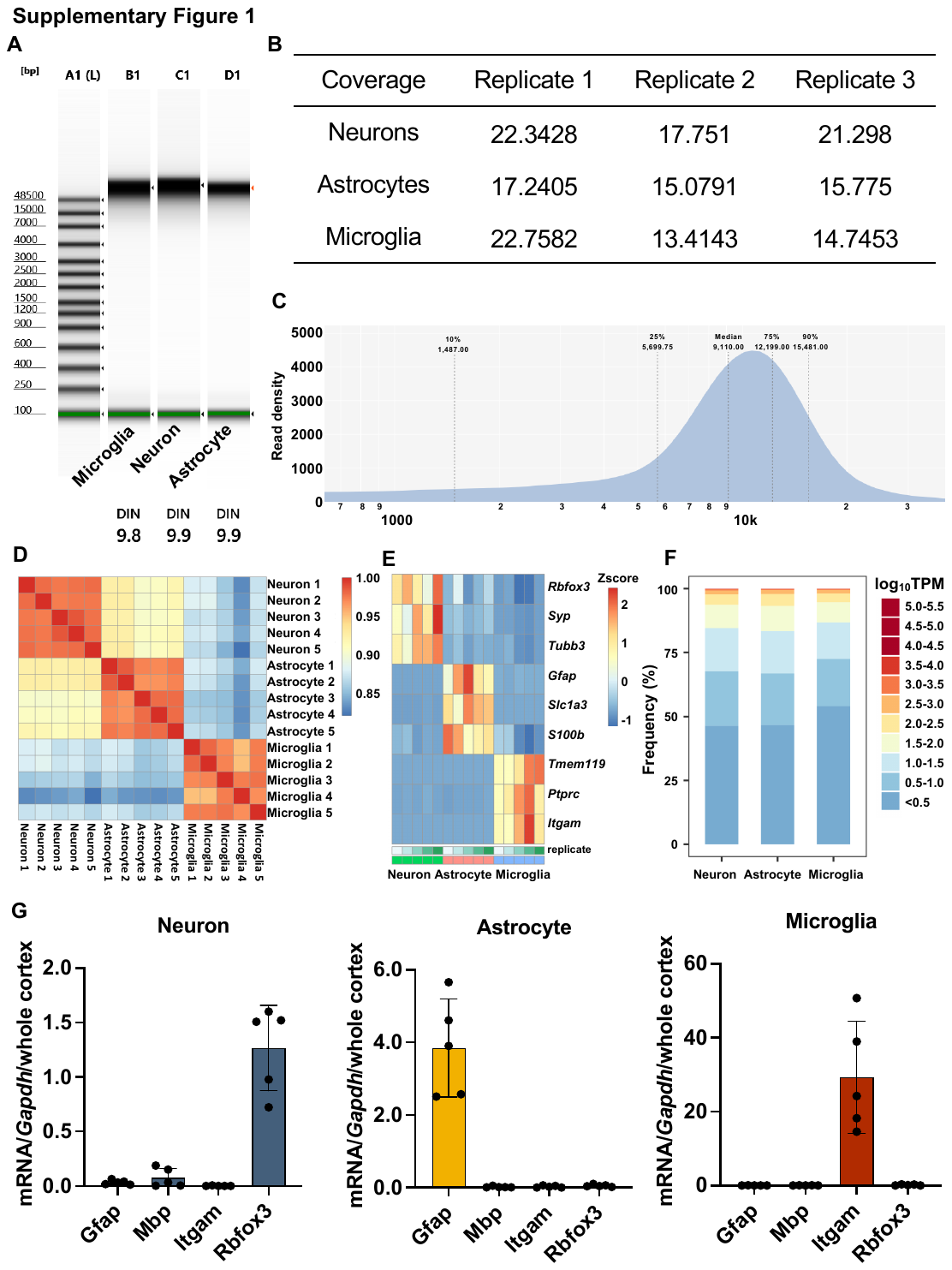

**Supplementary Figure 1. Nanopore sequencing and RNA sequencing quality check. A**, Tapestation results of HMW DNA used in Nanopore Sequencing. DNA integrity number (DIN) ranges from 10 (intact) to 1 (totally degraded). **B**, Average coverage of Nanopore libraries. **C**, Length distribution of sequencing reads. The median sequencing read length is 10kb. **D**, Correlation analysis for RNAseq libraries. Red means a high correlation score while blue means a relatively low correlation score. **E**, heatmap of the expression level of cell markers in different libraries. **F**, Gene expression level distribution of astrocytes, neurons and microglia. Log10 TPM was used to show the mRNA expression level. **G**, qPCR results of the isolated neurons, astrocytes and microglia. qPCR results demonstrate relative mRNA expression of gene glial fibrillary acidic protein (*Gfap*, marker for astrocyte), myelin basic protein (*Mbp*, marker for oligodendrocytes), integrin alpha M chain (*Itgam*, marker for microglia) and RNA binding fox-1 homolog 3 (*Rbfox3*, marker for neurons) in neurons, astrocytes and microglia.

**Supplementary Figure 2.** Differentially (hydroxy)methylated sites. **A** and **C**, Distribution of hypermethylated DMS between astrocytes and microglia (**A**) and between neuron and microglia (**C**). 8,697,812 DMSs were identified between astrocytes and microglia while 4,729,928 DMSs were identified between neuron and microglia. For astrocytes and microglia comparison, 89.2% of DMSs are hypermethylated in microglia (7,761,424 DMSs) while 10.8% of DMSs are hypermethylated in astrocytes (936,388 DMSs). From the comparison between neurons and microglia, we identify 83.5% of DMSs are hypermethylated in microglia (3,949,509 DMSs) while 16.5% of DMSs are hypermethylated in neurons (780,419 DMSs). **B** and **D**, DMSs between astrocytes and microglia (**B**) and between neuron and microglia (**D**). All the CpG sites are plotted based on their chromosome location and 5mC level difference between astrocytes and microglia (astrocytes served as the control) or 5mC level difference between neuron and microglia (neurons served as the control). **E** and **G**, Distribution of hypermethylated DhMS between astrocytes and microglia (**E**) and between neuron and microglia (**G**). 6,493,806 DhMS were identified between astrocytes and microglia while 845,004 DhMSs were identified between neuron and microglia. About 99.95% of DhMSs are hypermethylated in astrocytes (6,490,695 DhMSs) and 98.8% of DhMSs are hypermethylated in neurons (834,489 DhMSs) compared to microglia. **F** and **H**. DhMSs between astrocytes and microglia (**F**) and between neuron and microglia (**H**). All the CpG sites are plotted based on their chromosome location and 5hmC level difference between astrocytes and microglia (astrocytes served as the control) or between neuron and microglia (neurons served as the control).

**Supplementary Figure 3.** Differentially methylated regions. **A, C** and **E**, Distribution of hypermethylated differentially methylated regions between astrocytes and neurons (**A**), astrocytes and microglia (**C**) and neuron and microglia (**E**). In total, 509,214 DMRs were identified between astrocytes and neurons. 660,151 DMRs were identified between astrocytes and microglia while 485,234 DMRs were identified between neurons and microglia. Compared to neurons and microglia, the majority of DMRs are hypomethylated in astrocytes (compared to neurons: 65.2%/331,858 DMRs, compared to microglia: 78.5%/518,425 DMRs). In the comparison between neurons and microglia, 17.1% DMRs (83,183 DMRs) are hypermethylated in the neurons. **B, D** and **F**. DMRs between astrocytes and neurons (astrocytes served as the control) (**B**), astrocytes and microglia (astrocytes served as the control) (**D**) and neuron and microglia (neurons served as the control) (**F**). All the regions are plotted based on their chromosome location and 5mC level difference.

**Supplementary Figure 4.** Differentially hydroxymethylated regions. **A, C** and **E**, Distribution of hypermethylated differentially hydroxymethylated regions between astrocytes and neurons (**A**), astrocytes and microglia (**C**) and neuron and microglia (**E**). In total, 389,629 DhMRs were identified between astrocytes and neurons. 498,948 DhMRs were identified between astrocytes and microglia while 150,002 DhMRs were identified between neurons and microglia. Almost all the DhMRs are hypermethylated in astrocytes (compared to neurons: 97.6%/381,254 DhMRs, compared to microglia: 99.6%/496,813 DhMRs). In the comparison between neurons and microglia, 95.4% DhMRs (143,053 DhMRs) are hypermethylated in neurons. **B, D** and **F**, DhMRs between astrocytes and neurons (astrocytes served as the control) (**B**), astrocytes and microglia (astrocytes served as the control) (**D**) and neuron and microglia (neurons served as the control) (**F**). All the regions are plotted based on their chromosome location and 5hmC level difference.

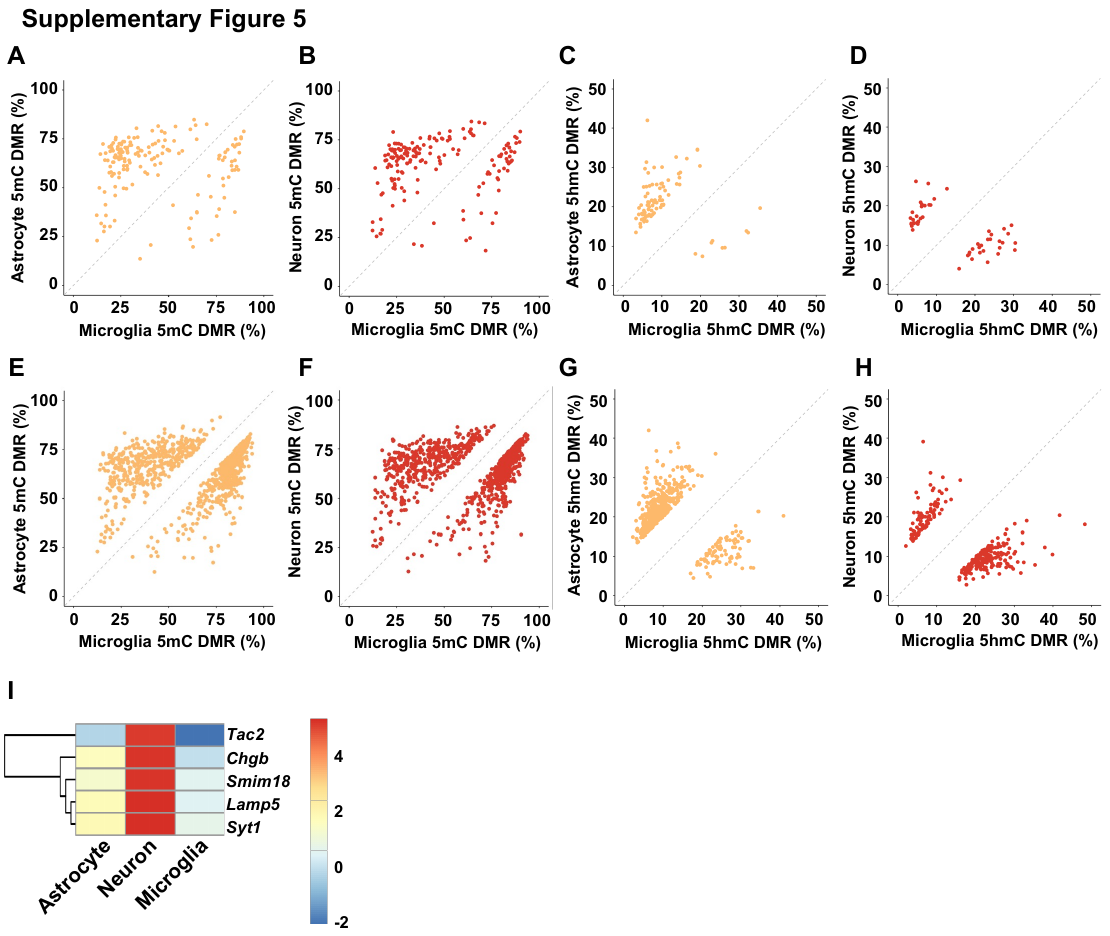

**Supplementary Figure 5. 5mC and 5hmC sites methylation level in the promoter region of microglia specific genes. A-B**, Methylation level of 5mC DMRs between astrocytes and microglia (**A**) and neurons and microglia (**B**) in the promoter region of microglia specific genes. **C-D**, Methylation level of 5hmC DMRs between astrocytes and microglia (**C**) and neurons and microglia (**D**) in the promoter region of microglia specific genes. **E-F**, Methylation level of 5mC DMRs between astrocytes and microglia (**E**) and neurons and microglia (**F**) in the gene body of microglia specific genes. **G-H**, Methylation level of 5hmC DMRs between astrocytes and microglia (**G**) and neurons and microglia (**H**) in the gene body of microglia specific genes. **I**, Gene expression level of potential neuron specific genes in astrocytes, neurons and microglia.
