## Supplemental Table 1 for "Decoding the Epigenetic Landscape: Insights into 5mC and 5hmC Patterns in Mouse Cortical Cell Types"

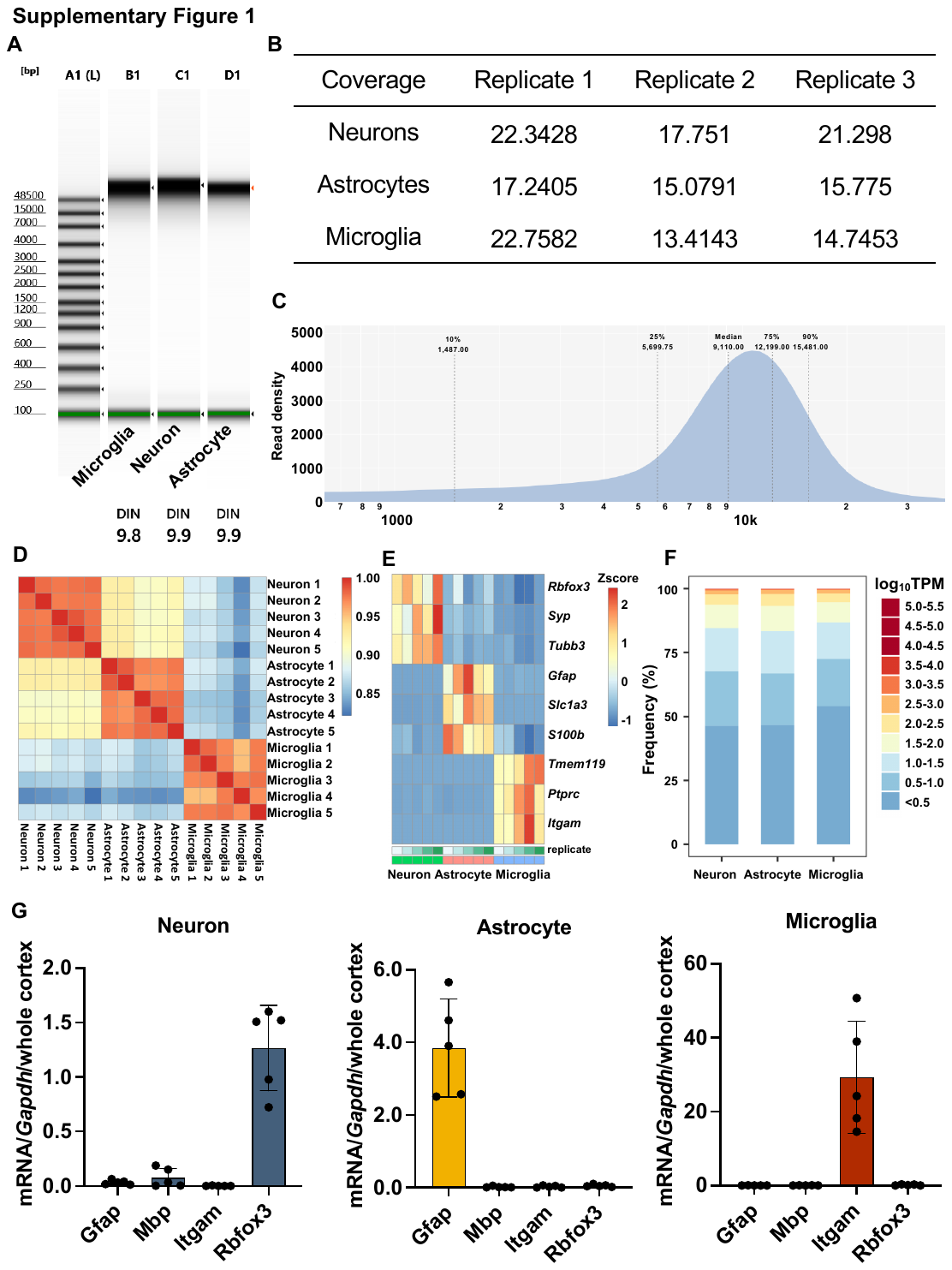

**Supplementary Figure 1. Nanopore sequencing and RNA sequencing quality check. A**, Tapestation results of HMW DNA used in Nanopore Sequencing. DNA integrity number (DIN) ranges from 10 (intact) to 1 (totally degraded). **B**, Average coverage of Nanopore libraries. **C**, Length distribution of sequencing reads. The median sequencing read length is 10kb. **D**, Correlation analysis for RNAseq libraries. Red means a high correlation score while blue means a relatively low correlation score. **E**, heatmap of the expression level of cell markers in different libraries. **F**, Gene expression level distribution of astrocytes, neurons and microglia. Log10 TPM was used to show the mRNA expression level. **G**, qPCR results of the isolated neurons, astrocytes and microglia. qPCR results demonstrate relative mRNA expression of gene glial fibrillary acidic protein (*Gfap*, marker for astrocyte), myelin basic protein (*Mbp*, marker for oligodendrocytes), integrin alpha M chain (*Itgam*, marker for microglia) and RNA binding fox-1 homolog 3 (*Rbfox3*, marker for neurons) in neurons, astrocytes and microglia.

**Supplementary Figure 4.** Differentially hydroxymethylated regions. **A, C** and **E**, Distribution of hypermethylated differentially hydroxymethylated regions between astrocytes and neurons (**A**), astrocytes and microglia (**C**) and neuron and microglia (**E**). In total, 389,629 DhMRs were identified between astrocytes and neurons. 498,948 DhMRs were identified between astrocytes and microglia while 150,002 DhMRs were identified between neurons and microglia. Almost all the DhMRs are hypermethylated in astrocytes (compared to neurons: 97.6%/381,254 DhMRs, compared to microglia: 99.6%/496,813 DhMRs). In the comparison between neurons and microglia, 95.4% DhMRs (143,053 DhMRs) are hypermethylated in neurons. **B, D** and **F**, DhMRs between astrocytes and neurons (astrocytes served as the control) (**B**), astrocytes and microglia (astrocytes served as the control) (**D**) and neuron and microglia (neurons served as the control) (**F**). All the regions are plotted based on their chromosome location and 5hmC level difference.

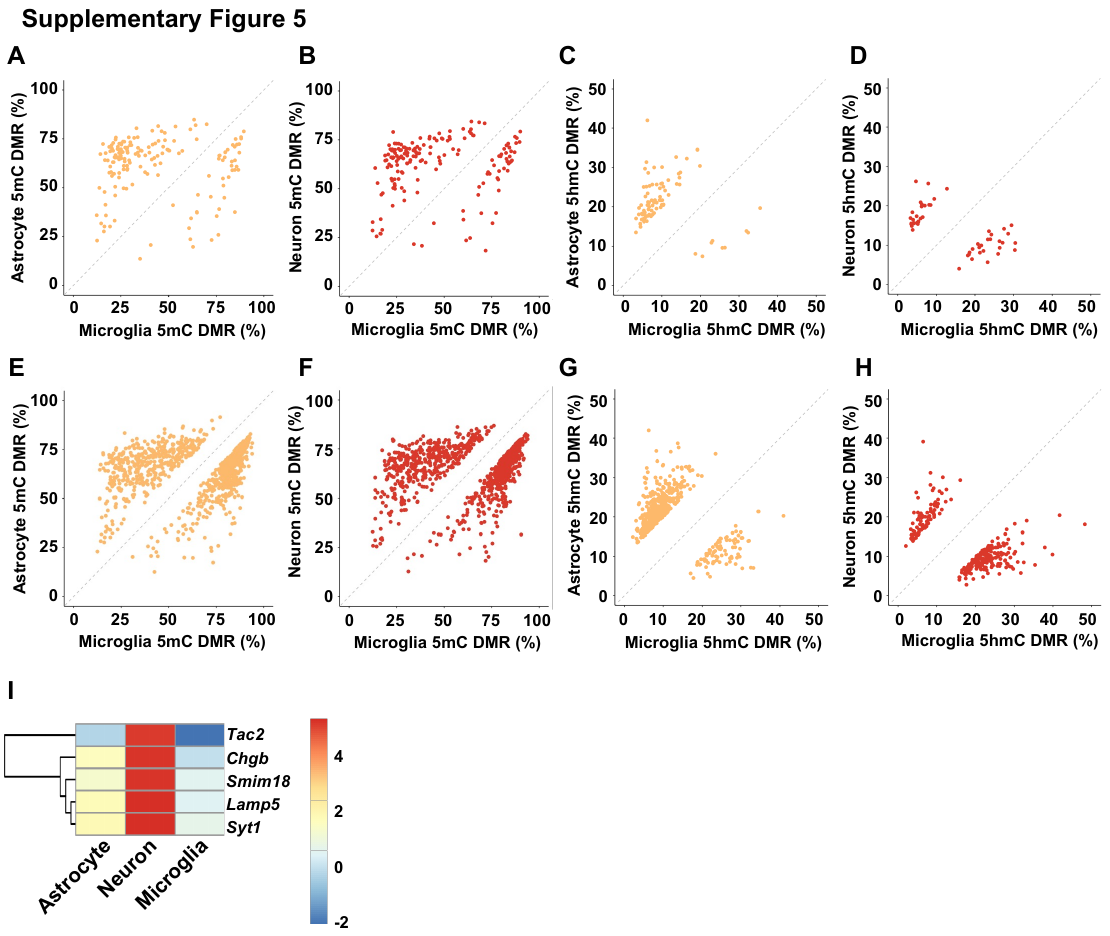

**Supplementary Figure 5. 5mC and 5hmC sites methylation level in the promoter region of microglia specific genes. A-B**, Methylation level of 5mC DMRs between astrocytes and microglia (**A**) and neurons and microglia (**B**) in the promoter region of microglia specific genes. **C-D**, Methylation level of 5hmC DMRs between astrocytes and microglia (**C**) and neurons and microglia (**D**) in the promoter region of microglia specific genes. **E-F**, Methylation level of 5mC DMRs between astrocytes and microglia (**E**) and neurons and microglia (**F**) in the gene body of microglia specific genes. **G-H**, Methylation level of 5hmC DMRs between astrocytes and microglia (**G**) and neurons and microglia (**H**) in the gene body of microglia specific genes. **I**, Gene expression level of potential neuron specific genes in astrocytes, neurons and microglia.

**Supplementary Table 1. Cell Type specific genes.**

| Cell type | Genes |
| --- | --- |
| Astrocyte | Aldh1l1, Btbd17, Cbs, Chrdl1, Cideb, Cth, Cyp4f14, Cyp4f15, Entpd2, Ephx2, Fgfr3, Fjx1, Gabrg1, Gpc5, Hapln1, Myorg, Nat8f5, Ndp, Paqr6, Pdzph1, Phkg1, Plcd4, Ppp1r3g, Sox9 |
| Microglia | Abi3, Adgre1, Apbb1ip, BC035044, Bcl2a1a, Bcl2a1b, Bin2, Blnk, C5ar1, Ccl2, Ccl3, Ccl4, Ccl6, Ccl8, Ccl9, Ccr5, Cd14, Cd300c2, Cd33, Cd37, Cd48, Cd53, Cd84, Cd86, Clec4a2, Clec4a3, Clec5a, Cryba4, Csf1r, Csf3r, Cx3cr1, Cysltr1, Cyth4, Ear2, Fcgr1, Fcgr2b, Fermt3, Fgd2, Folr2, Fyb, Gna15, Gp9, Gpr183, Gpr34, Gpr84, H2-Oa, Havcr2, Hck, Hpgd, Hpgds, Hspb3, Il10ra, Il1a, Irf5, Irf8, Itgam, Itgb2, Lat2, Lcp2, Lpcat2, Lpxn, Mafb, Ms4a4a, Ms4a6b, Ms4a6c, Ms4a6d, Ms4a7, Ncf1, Ncf4, P2ry13, P2ry6, Pld4, Ppfia4, Retnla, Retnlg, Rgs1, Rhoh, Rnase6, Samsn1, Siglech, Sla, Slamf9, Slc11a1, Slc25a45, Slc2a5, Slc7a7, Spi1, Susd3, Tbxas1, Tgfbr1, Tifab, Tlr7, Tmem119, Tmem273, Tnfaip8l2, Tnfrsf13b, Tnfrsf17, Trem2, Upk1b, Vav1 |
